# Real-time genomic surveillance during the 2021 re-emergence of the yellow fever virus in Rio Grande do Sul State, Brazil

**DOI:** 10.1101/2021.07.16.452727

**Authors:** M.S. Andrade, F.S. Campos, A.A.S. Campos, F.V.S. Abreu, F.L. Melo, J.C. Cardoso, E. Santos, L.C. Born, C.M.D. Silva, N.F.D. Müller, C.H. Oliveira, A.J.J. Silva, D. Simonini-Teixeira, S. Bernal-Valle, M.A. Mares-Guia, G.R. Albuquerque, A.P. Sevá, A.P.M. Romano, A.C. Franco, B.M. Ribeiro, P.M. Roehe, M.A.B. Almeida

## Abstract

The yellow fever virus (YFV) re-emergence in Rio Grande do Sul, Brazil, raised big concerns and led the state to declare a Public Health Emergency of State Importance. Here, we generated near-complete genomes from the ongoing outbreak in Southern Brazil, aiming to better understand the phylogenetic aspects and also spatio-temporal dynamics of the virus. Our findings highlight the path and dispersion in Rio Grande do Sul and that YFV was reintroduced from São Paulo to the Rio Grande do Sul state through Paraná and Santa Catarina states, at the end of 2020.

## 1. INTRODUCTION

Yellow fever (YF) is a viral hemorrhagic fever caused by the Yellow Fever Virus (YFV), the prototype of the genus *Flavivirus*, family *Flaviviridae* (Monath, 2001). In South America, YFV is widely spread and maintained in a sylvatic cycle by transmission between non-human primates (NHP’s) and blood-feeding mosquitoes mainly from the genera *Haemagogus* (Monath and Vasconcelos 2015).

In Brazil, the area of YFV occurrence extends from the Amazon region, northern Brazil, to the state of Rio Grande do Sul (RS), the southernmost region of the country. The latter is sporadically affected when the virus overflows from the North (endemic area) to the Southeast and South Regions (Almeida et al., 2012; Delatorre et al., 2019; Figueiredo et al., 2020; Monath and Vasconcelos, 2015; Possas et al., 2018; Vasconcelos et al. 2004). During the YFV spreads, NHP’s deaths (hereafter called epizootics) precedes human cases, highlighting the importance of the epizootic surveillance system as a tool for early detection of YFV circulation. Prompt action in such episodes allows to actively increase vaccination coverage of human populations in the vicinity of the epizootic event as well as for mapping virus dispersion (Almeida et al., 2014; Romano et al., 2014).

Among South American NHP’s genera, *Alouatta* is particularly important for YF surveillance, because it is the most severely affected by YFV (Almeida et al., 2014; Hill et al., 2020; Mares-Guia et al., 2020; Moreno et al., 2013). In 2001-2002, following epizootics affecting *Alouatta* sp., a YF surveillance based on monitoring of living and dead NHP’s was implemented in RS. Unfortunately, between 2008-2009 a new outbreak occurred in RS, causing thousands of NHP’s deaths, and 21 human cases. In the subsequent twelve years of continued surveillance, no virus circulation was evidenced in the South Region (Almeida et al., 2012; 2014; 2019). Meanwhile, between 2014 - 2019, a new YFV foci was first detected in Tocantins state, in the North Region, and reached Southeast and South Regions, including Brazilian coast states Espírito Santo, Rio de Janeiro and São Paulo, causing the largest sylvatic outbreak ever recorded (Abreu et al., 2019b; Bonaldo et al., 2017; Brasil, 2014; Brasil, 2015; Brasil, 2017; Brasil, 2019; Cunha et al., 2019; Silva et al., 2020). Following these events, from 2019 onwards, YFV continued to spread towards the South Region, arriving in the southern states of Paraná and Santa Catarina and causing human cases and hundreds of epizootics (Brasil, 2019; DIVE, 2021; SESA, 2021).

Altogether, that YF outbreak affected ten states and the Federal District outside the Amazon and left behind about two thousand human cases and countless NHP’s deaths. That outbreak was monitored by real-time genomic surveillance (Brasil, 2019), and the phylogenetic analysis revealed the existence of at least two main viral sub-lineages occurring in Brazil in that period - the “Yellow Fever Virus Minas Gerais/São Paulo” (hereafter YFVMG/SP) and the “Yellow Fever Virus Minas Gerais/Espírito Santo/Rio de Janeiro” (hereafter YFVMG/ES/RJ) (Delatorre et. al., 2019).

Since the virus entered the RS neighboring state (Santa Catarina) in 2019 (Brasil, 2019; DIVE, 2020), following its progression to the south, health authorities of RS focused its surveillance efforts at border municipalities. Aimed for this, a surveillance team from the State Health Department was working to promote vaccination and raise awareness of people who live in that region to report NHP’s deaths (CEVS, 2021). Noteworthy, at the end of 2020 and beginning 2021, the virus continued its route southwards causing the first cases of NHP’s (*Alouatta guariba clamitans*) deaths on RS.

Although the virus has crossed Paraná and Santa Catarina territories, to date, no YFV genomes of viruses circulating in these states were publicly available and little is known about the dynamics of spatial spread of the YFV in southern Brazil. In this way, aiming to fill this information gap, here we sequence the first YFV genomes from the edge of the current YFV expansion wave in the South Region of Brazil, recovered from NHP’s samples during YF epizootics surveillance in RS.

## 2. METHODS

### 2.1 Ethics statement

This study comprised analysis of routinely collected surveillance data performed by the state and municipalities health departments and followed guidelines of the Ministry of Health of Brazil and Brazilian National Committee for Ethics in Research. All samples were obtained from dead NHPs. This study followed the guide to epizootic surveillance in non-human primates and entomology applied to yellow fever surveillance (Brasil, 2017) and Institutional Animal Care and Use Committees (IACUCs) review the Use of Nonhuman Primates (Tardif et al., 2013). This study was conducted in accordance with Brazilian legislation under the SISBIO/ICMBio/MMA authorizations for activities with scientific purpose 75734-1 and SISGEN license AF40BCA.

### 2.2 Sample collection

All NHP’s samples were collected in municipalities in the Northeast Region of the state of Rio Grande do Sul, Brazil. Samples from liver, kidney, lung, spleen and heart were collected in the field from dead animals and were kept refrigerated (4°C) or frozen in dry ice and dispatched to the central office of the State Health Department, where they were stored at - 80°C. All collections followed biosafety protocols and were in accordance with the state YF surveillance strategy carried out by the Environmental Health Surveillance Division, State Center of Health Surveillance, State Health Department of RS, and the Ministry of Health of Brazil. Data concerning the geo-located origin of the animals, date of sampling and post-mortem findings were registered.

### 2.3 RT-qPCR

YFV RNAs were extracted from NHP’s tissue (liver and kidney) samples spotted on FTA^®^ classic filter paper (Whatman). A hole punch was used to excise a single dried 6 mm diameter circle. To avoid carryover contamination, the instrument was disinfected with bleach, water and ethanol 100% (Bonne et al., 2008). The material was then lysed using proteinase K and lysis buffer containing carrier RNA at 56 °C, for 15 min, according to PureLink Viral Mini kit protocol (Invitrogen). The lysates samples were finally extracted using automated extraction (Loccus, Extracta Kit FAST). Viral RNA was detected using two previously published RT-qPCR protocols (Domingo et al., 2012). Samples which were positive at RT-qPCR with cycle thresholds (CT) below 25 were sent to Baculovirus Laboratory, in University of Brasilia, for sequencing.

### 2.4 Genome sequencing

All samples that met the previous criteria were submitted to cDNA synthesis protocol using LunaScript™ RT SuperMix Kit (NEB) following the manufacturer’s instructions. Then, a multiplex tiling PCR was attempted using the previously published YFV primers (Faria et al., 2018) and 40 cycles (denaturation: 95°C/15 s and annealing/extension: 65°C/5 min) of PCR using Q5 high-fidelity DNA polymerase (NEB). Amplicons were purified using 1× AMPure XP beads (Beckman Coulter), and cleaned-up PCR product concentrations were measured using a QuantiFluor® dsDNA System assay kit on a Quantus™ Fluorometer (Promega). DNA library preparation was performed using the Ligation sequencing kit SQK-LSK309 (Oxford Nanopore Technologies) and the Native barcoding kit (EXP-NBD104 and EXP-NBD114; Oxford Nanopore Technologies, Oxford, UK). The sequencing library (23 samples and a negative control per run) was loaded onto a R9.4 flow cell (Oxford Nanopore Technologies) and sequenced between 6 to 18 hours using MiNKOW software. The RAMPART (Version 1.2.0, ARTIC Network) package was used to monitor coverage depth and genome completion. The resulting Fast5 files were basecalled and demultiplexed using Guppy (Version 4.4.2, Oxford Nanopore Technologies). Variant calling and consensus genome assembly was carried out with Medaka (Version 1.0.3, Oxford Nanopore Technologies) using the sequence JF912190 as the reference genome.

### 2.5 Phylogenetic analysis

To perform phylogenetic analysis, we selected from NCBI all near-complete YFV sequences (YFV-set-1, n=359, sequences > 8 kb excluding sequences from vaccine and patents). Then, we make a subset of sequences belonging to recent (2015 to 2021) extra-Amazonian region waves, including clades YFV_MG/ES/RJ_ and YFV_MG/SP_ (YFV-subset, n=264). Metadata as samples collection date and geographic coordinates were retrieved from GenBank files or from genome associated publications (manual curation). Genomes generated here (n = 23) combined with YFV-set-1 were aligned with MAFFT v.7.480 (Katoh and Standley, 2013). The Maximum-likelihood tree was inferred using IQTREE, with the GTR+F+I+┌_4_ model. YFV-subsets combined with the newly determined genomes were used to construct a time-scaled tree in Nextstrain (https://nextstrain.org/ncov). The new genome sequences will be sent to the NCBI GenBank database to obtain accession numbers.

### 2.6 Epidemiological and geographic information

Epidemiological data about human cases (from 2016 to March 8^th^, 2021) and epizootics (from 2014 to March 8^th^, 2021) related to YF in Brazil, was provided by the Ministry of Health of Brazil by the law of access to information. Numbers of Epizootics in RS were provided by Division of Environmental Health Surveillance from Rio Grande do Sul State Health Department. Maps presenting the results were generated through free software QGIS version 2.18.

## 3. RESULTS

Between January and March 08^th^ (date of collection of our last sequenced sample), 78 epizootics were notified to RS State Health Department and Ministry of Health of Brazil surveillance, of which, in 42 (54%) sample collection could be performed. From these, 34 (81%) tested positive for YFV by RT-qPCR assay. All confirmed YFV-positive NHP’s were *Alouatta guariba clamitans* and it became evident that, at the date of collection of our last sequenced sample, the virus remained in circulation in the state for at least 10 weeks, with epizootics occurring mainly in the Northeast Region of RS near to the border with Santa Catarina state (**Figure 1**).

**Figure 1.**
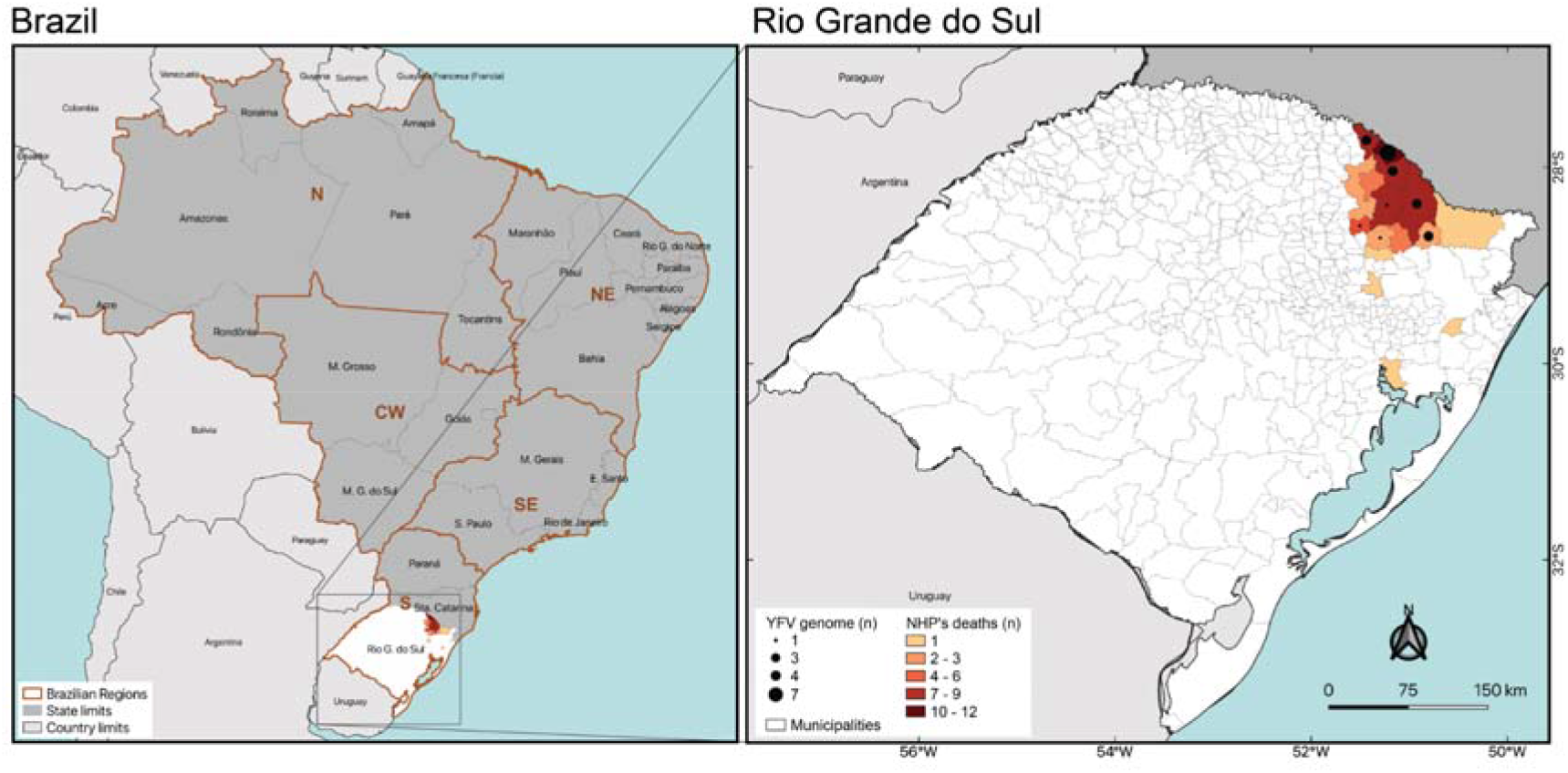
Geographical distribution of YFV NHP cases in Rio Grande do Sul. The municipalities with NHP’s deaths positive at RT-qPCR are highlighted and genome numbers per municipality are shown.

Twenty-three near-complete YFV genomes were generated from liver samples from dead NHP’s collected in eight municipalities of RS. At RT-qPCR, sequenced samples displayed median Ct values of 12 (range 8 - 20) (**Table 1**). Sequences generated here from RS NHP’s samples clustered within the South America I clade (Mir et al., 2017; Souza et al., 2010) (**Figure 2**) and did not group with previous genomes from RS, JF912189 (2001) and KY861728 (2008) (Vasconcelos et al., 2003) indicating a re-emergence of YFV in RS.

**Table 1.** List of genomes generated in this study showing date of collection, sample name, latitude, longitude, cycle threshold (Ct) and coverage.

| Date | Sample name | Lat | Long | Ct | Coverage |
| --- | --- | --- | --- | --- | --- |
| 25 January | Pinhal da Serra 01 | -27.8757 | -51.2260 | 20 | 194 |
| 03 February | Pinhal da Serra 02 | -27.8757 | -51.2260 | 11 | 156 |
| 03 February | Pinhal da Serra 03 | -27.8757 | -51.2260 | 11 | 135 |
| 08 February | Monte Alegre dos Campos 01 | -28.5355 | -51.5023 | 10 | 30 |
| 08 February | Pinhal da Serra 05 | -27.8346 | -51.1995 | 10 | 284 |
| 11 February | Pinhal da Serra 07 | -27.8843 | -51.1632 | 13 | 53 |
| 19 February | Esmeralda 01 | -28.0930 | -51.1124 | 12 | 360 |
| 19 February | Pinhal da Serra 08 | -27.8933 | -51.1116 | 9 | 303 |
| 19 February | Pinhal da Serra 09 | -27.8403 | -51.2505 | 10 | 1572 |
| 20 February | Muitos Capões 01 | -28.2196 | -51.2135 | 8 | 309 |
| 22 February | Barracão 02 | -27.7315 | -51.3688 | 11 | 247 |
| 22 February | Barracão 03 | -27.7315 | -51.3688 | 9 | 153 |
| 22 February | Barracão 04 | -27.7315 | -51.3688 | 15 | 666 |
| 22 February | Vacaria 01 | -28.2900 | -50.8116 | 9 | 568 |
| 22 February | Vacaria 02 | -28.2900 | -50.8116 | 10 | 416 |
| 24 February | Esmeralda 02 | -28.1698 | -50.9228 | 15 | 207 |
| 24 February | Vacaria 04 | -27.9216 | -51.2187 | 11 | 533 |
| 25 February | Esmeralda 03 | -27.9754 | -51.0557 | 9 | 153 |
| 25 February | Esmeralda 04 | -27.9754 | -51.0557 | 11 | 294 |
| 26 February | Monte Alegre dos Campos 02 | -28.7591 | -51.2527 | 11 | 84 |
| 03 March | Monte Alegre dos Campos 03 | -28.6724 | -50.7997 | 14 | 722 |
| 04 March | André da Rocha 02 | -28.5850 | -51.5757 | 20 | 1008 |
| 08 March | Ipê 01 | -28.7591 | -51.2527 | 12 | 2024 |

**Figure 2.**
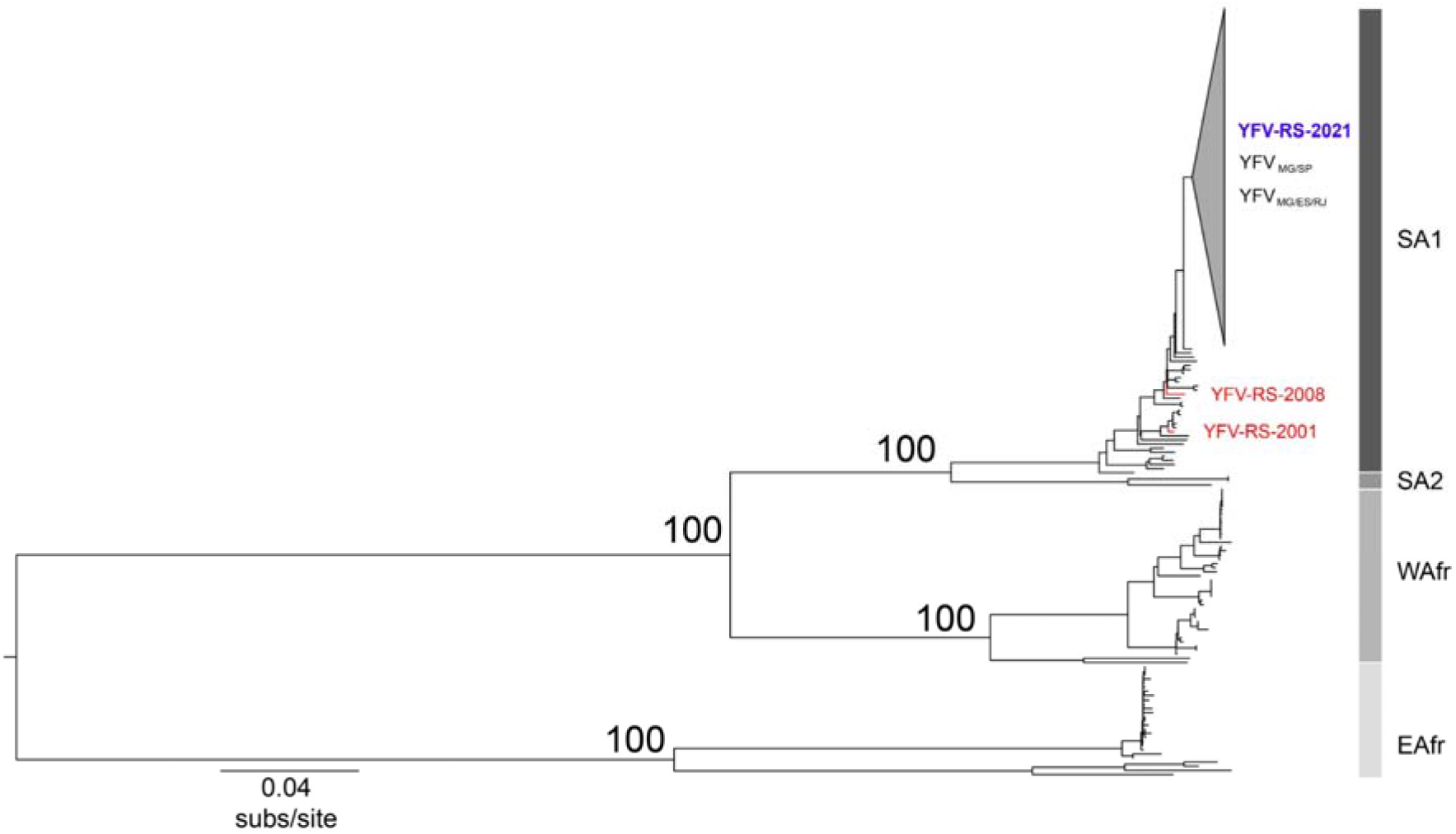
Phylogenetic tree of YFV based on 381 near-complete genomes. The gray collapsed group includes YFV_MG/ES/RJ_ and YFV_MG/SP_ clades and all genomes from RS sequenced in this study. South America I, South America 2, West Africa and East Africa genotypes are indicated. Previous genomes from RS, JF912189 (2001) and KY861728 (2008), are highlighted in red.

To understand the spatial and evolutionary dynamics of recent YFV dispersion from endemic areas to southern regions of Brazil we made an analysis using a subset of sequences belonging to the South America 1 genotype on Nextstrain. The time-scaled phylogenetic tree (**Figure 3A**) shows that YFV from RS sequenced here are clustered with the YFV_MG/SP_ sub-lineage, named Yellow Fever Virus Minas Gerais/São Paulo/Rio Grande do Sul (hereafter YFV_MG/SP/RS_), revealing that the origin of this isolates is São Paulo. Furthermore, the time-scaled phylogenetic tree suggests more than one entry in RS. Despite the lack of genomic data from Paraná and Santa Catarina, epidemiological data of epizootics and human cases due to YF from São Paulo and South Region (Paraná, Santa Catarina and RS), as shown in **Figure 3B**, suggest that the recent YFV dispersion wave achieve RS passing through Paraná (2018/2020) and Santa Catarina (2019/2021).

**Figure 3.**
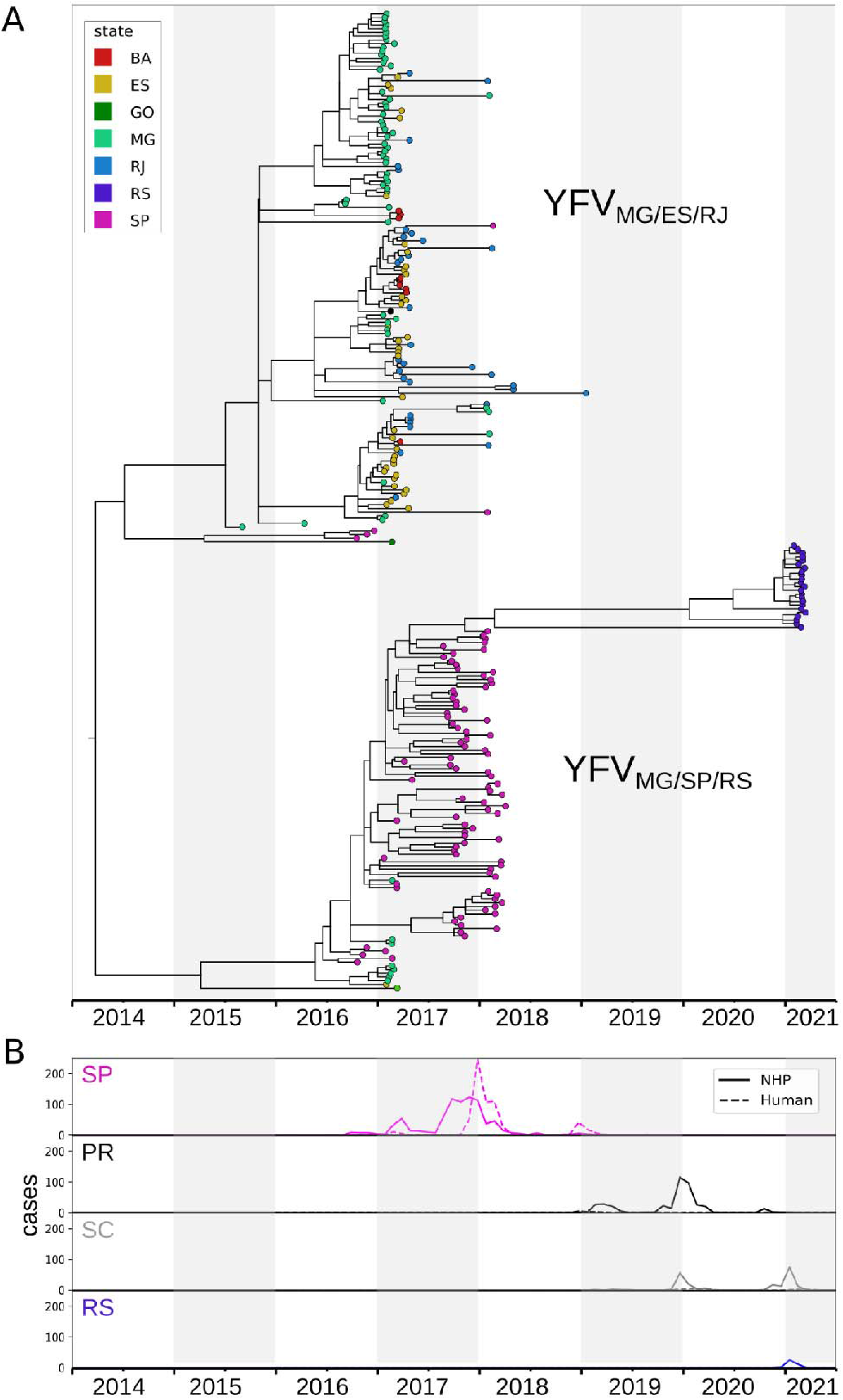
Spatio-temporal YFV spread. A) Time-scaled phylogenetic tree of YFV_MG/ES/RJ_ and YFV_MG/SP/RS_ sub-lineages. B) Epizootics and human cases due to YF, reported in São Paulo (SP), Paraná (PR), Santa Catarina (SC) and Rio Grande do Sul (RS) by month (Source: Ministry of Health of Brazil).

## 4. DISCUSSION

In this study we applied real-time genomic surveillance to obtain sequences of YFV from NHP’s found dead by the YF surveillance system in RS state, southern Brazil, at the beginning of the re-emergence of the virus in 2021. The new introduction of YFV in RS has raised big concerns and led the state to declare a Public Health Emergency of State Importance. To better understand YFV genetic distribution during the recent outbreak in Southern Brazil, we generated 23 complete and nearly complete genomes from the first peak of the epidemic curve from NHP’s cases in the northeast of RS state.

In Brazil, over the past five years, previous genetic analyses have indicated the circulation of at least two distinct sub-lineages, in two different transmission routes. The first one was YFV MG/ES/RJ, which circulated in the east of Minas Gerais in 2016, entering the west of Espírito Santo and following to the north of Rio de Janeiro. From there, in 2017, the virus spread southwards, arriving at the border with São Paulo in 2018 (Bonaldo et al., 2017; Delatorre et al., 2019; Faria et al., 2018; Goméz et al, 2018). About march 2017, NHP’s samples infected by this sub-lineage were also found in Bahia, a Northeast Region state in Brazil (Goes de Jesus et al., 2020). Meanwhile, the second sub-lineage, named YFV_MG/SP_, circulated in the west of Minas Gerais and the northwest of São Paulo in 2016-2017, from where it advanced towards the south and east of that state, reaching the most densely populated region of the country between 2017-2018 (Hill et al., 2020; Rezende et. al., 2018). From 2018 to 2021 YFV was detected in NHP and humans in Paraná and/or Santa Catarina (Brasil, 2019; DIVE, 2021; SESA, 2021). Although the lack of genomic information available from Paraná and Santa Catarina, the spatiotemporal distribution of epizootic and human cases in South Region combined with phylogenetic analysis presented here suggest that YFV wave toward South Region of Brazil is a continuity of sub-lineage YFV_MG/SP_ (now called YFV_MG/SP/RS_) expansion.

Previous records of occurrence of YF in RS reveal that the border between Argentina and Brazil (northwest of RS) is historically the first affected area, as happened in 1947, 1966, 2001, 2002 and 2008/2009 (Almeida et al. 2012, 2014; Bejarano, 1974; Cardoso et al., 2010; Franco, 1969; Vasconcelos et al., 2003). In the epizootic here reported, the virus followed a different route reaching northeast of RS, a route which matches with recent dispersion of YFV in Santa Catarina and Paraná, coming from the southeast of São Paulo state. Furthermore, our data lead us to believe that samples from Santa Catarina and Paraná, if sequenced, might be grouped with sub-lineage YFV_MG/SP/RS_. Sequencing samples from these states are essential to better understand the dispersion pattern of YFV while circulating in the South Brazil Region.

Interestingly, the sequences obtained here, represent virus samples of the farthest place reached by YFV dispersion in Brazil, since it spread from the endemic area, about 2014, in this expansion wave. Rio Grande do Sul is the south limit for NHP’s distribution in the Americas (Printes and Liesenfeld, 2001) and the farthest state from YF endemic area, which is the source of YFV in all extra-Amazon circulations (Monath and Vasconcelos, 2015). Consequently, it is expected to be the south limit for YFV spread in the Americas too. Possibly, the 2021 YFV lineage from RS presents accumulated genetic changes over seven years of circulation, since the first detection of an extra endemic area, in 2014, in the State of Tocantins (Brazil, 2019).

Epidemic outbreaks in the sylvatic cycle of YFV demands a high density of competent vectors and susceptible NHP’s presence, acting as an amplifier for the virus (Abreu et al., 2019b; Cardoso et al., 2010; Mares-Guia et al., 2020; Pinheiro et al., 2019). Importantly, the YFV expansion wave (2014-2021) revealed the suitability of climate and ecological conditions for the occurrence of YFV outbreaks in several Brazilian regions, even though in the three southernmost states of Brazil (Paraná, Santa Catarina and RS), which have a subtropical temperate climate and great loss of forest areas, through which viral circulation can occur (Abreu et al., 2019a; Delatorre et al., 2019; Sacchetto et al., 2020a; Sacchetto et al., 2020b; Rosa et al., 2021). Therefore, it is important to strengthen surveillance. Our findings confirm the need for continued surveillance for early detection of outbreaks and the use of genetic tools to determine the origin and understanding of YFV entry and dispersion in RS.

Ultimately, we present here the first sequences of the front of the current YFV wave heading south and show that these sequences are related with the lineage identified in SP during YFV circulation in that state in 2017-2018. However, further studies of the virus dispersion including ecological, climatic, and anthropogenic factors associated with the disease cases are needed to improve predictive power, allowing rapid decisions into surveillance and prevention efforts. Furthermore, we are performing continuous real-time genomic surveillance at RS, once there is an ongoing outbreak over NHP’s, and new genome sequencing and ecological analysis are in progress. Lastly, our work shows the importance of capacity building for inter-institutional exchange of data and human resources, to strengthen the epidemic surveillance and outbreak management during pandemics.

## Acknowledgments

We acknowledge the contributions of the Division of Environmental Health Surveillance from Rio Grande do Sul State Health Department for the important work over the years of epidemiological surveillance and sample collection. The authors would like to thank the effort of the RS Yellow Fever Surveillance Reference Team that was at the forefront of the preparation to face the arrival of the virus in the state as well as in field investigations. We are also grateful to countless colleagues from municipalities’ health departments, who conducted the investigation of epizootics collecting samples in the field and to the Ministry of Health’s Arbovirus Surveillance Team. The Yellow Fever Brazil project (Febre Amarela BR: https://www.febreamarelabr.com.br/) is supported by grants from Conselho Nacional de Desenvolvimento Científico e Tecnológico and Departamento de Ciência e Tecnologia of Secretaria de Ciência, Tecnologia e Insumos Estratégicos of Ministério da Saúde (CNPq/Decit/SCTIE/MS grant number 443215/2019-7). M.S.A. is granted a post-doctoral scholarship (DTI-A) from CNPq.

